# Neural signatures of dynamic emotion constructs in the human brain

**DOI:** 10.1101/200873

**Authors:** Tijl Grootswagers, Briana L. Kennedy, Steven B. Most, Thomas A. Carlson

**Author notes:** T. Grootswagers and B.L. Kennedy contributed equally to this work. Please address correspondence to: Briana L. Kennedy, University of Southern California, Or Tijl Gootswagers, University of Sydney.

## Abstract

How is emotion represented in the brain: is it categorical or along dimensions? In the present study, we applied multivariate pattern analysis (MVPA) to magnetoencephalography (MEG) to study the brain’s temporally unfolding representations of different emotion constructs. First, participants rated 525 images on the dimensions of valence and arousal and by intensity of discrete emotion categories (happiness, sadness, fear, disgust, and sadness). Thirteen new participants then viewed subsets of these images within an MEG scanner. We used Representational Similarity Analysis (RSA) to compare behavioral ratings to the unfolding neural representation of the stimuli in the brain. Ratings of valence and arousal explained significant proportions of the MEG data, even after corrections for low-level image properties. Additionally, behavioral ratings of the discrete emotions fear, disgust, and happiness significantly predicted early neural representations, whereas rating models of anger and sadness did not. Different emotion constructs also showed unique temporal signatures. Fear and disgust – both highly arousing and negative – were rapidly discriminated by the brain, but disgust was represented for an extended period of time relative to fear. Overall, our findings suggest that 1) dimensions of valence and arousal are quickly represented by the brain, as are some discrete emotions, and 2) different emotion constructs exhibit unique temporal dynamics. We discuss implications of these findings for theoretical understanding of emotion and for the interplay of discrete and dimensional aspects of emotional experience.

## 1. Introduction

Emotions are a potent part of our daily lives; the way in which we characterize and experience them is the focus of contentious and ongoing debate (see Russell, 2009). Modern neuroimaging and statistical techniques allow for new ways to examine how emotions are delineated in the brain and the indices provided by such techniques might shed light on the most appropriate ways to define or understand them (see Hamann, 2012; Kragel & LaBar, 2016 for reviews).

Two traditional, polar perspectives in the field suggest that emotions exist either as discrete entities or along dimensional space. The *discrete* emotion perspective suggests that a number of nominal, basic, specific emotions exist in categorical space (Ekman, 1992; Izard, 1992). In its most strict interpretation, the basic emotion perspective suggests that innate emotions comprise the emotional space as separate entities with unique and distinct physiological correlates: e.g., fear, sadness, anger, surprise, joy, contempt, disgust (Ekman, 1992). In contrast, the traditional *dimensional* perspective suggests that emotions exist along graded dimensions, such as valence (positive vs. negative) and arousal (activated vs. deactivated), and that the spectrum of emotional experience can be characterized by where they fall along these dimensions (Bradley & Lang, 1994; Rubin & Talarico, 2009; Russell, 1980). For example, in this framework, sadness exists within the quadrant of dimensional space where negative valence and low arousal intersect, whereas fear and disgust can be characterized by the intersection of negative valence with high arousal. More modern versions of the dimensional approach have included additional dimensions, such as approach-vs. withdrawal- related value (e.g., Harmon-Jones, Gable, & Price, 2013), or potency and unpredictability (Fontaine, Scherer, Roesch, & Ellsworth, 2007). Both perspectives have been supported through decades of research (Russell, 2009), resulting in little consensus about the underlying organizational structure of emotional constructs (Hamann, 2012).

Modern alternatives to these traditional theories of emotion include constructivist, network-based approaches (Barrett, 2017; Cunningham, Dunfield, & Stillman, 2014; Lindquist, Siegel, Quigley, & Barrett, 2013). For example, Barrett’s “theory of constructed emotion” suggests that emotions are experienced and learned based on previous, similar experiences, and derived from a network-based representation (Barrett, 2009, 2017). In this framework, seemingly discrete emotions are experienced by applying conceptual knowledge (often derived from previous experience and prediction processes) to interoceptive sensations that can often be characterized in terms of (but not limited to) valence and arousal dimensions (Barrett, 2017). Previous iterations of this model referred to the dimensional underpinnings as “core affect” (Russell & Barrett, 1999). Although constructivist accounts might be interpreted as the antithesis of a discrete emotions perspective, in some ways they also serve as a vehicle to reconcile discrete emotion- and dimensional- views. We might predict, for example, that individual ratings of valence and arousal (to the degree that such ratings have face value) account for a large degree of variance in the brain’s response to emotional stimuli, and that such dimensions should predict variance early in the temporal unfolding of such neural representations. We might also expect ratings of discrete emotions to account for some variance, but with the substance of their predictive value observed later in time.

Note that such a formulation raises additional questions: are there reliable differences in the rate at which representations of different discrete emotions are constructed? And, if it is correct that much of emotional experience is built on a foundation that includes dimensions of valence and arousal, is this necessarily true for all discretely experienced emotions? In the case of fear, for example, one influential suggestion has been that the amygdala (heavily implicated in fear) connects with other brain regions both via a cortical route and via routes that rapidly bypass cortical regions involved in attention and awareness (e.g., LeDoux, 2000; Morris, Scott, & Dolan, 1999; Tamietto & de Gelder, 2010). Although this suggestion has been challenged in recent years (Pessoa & Adolphs, 2010), the notion highlights the possibility that the neural representation of fear may unfold more rapidly than that of other emotions, perhaps so rapidly as not to be preceded by neural signatures of valence and arousal.

Multivariate pattern analysis (MVPA) or “brain decoding” techniques can be used to identify patterns of activation that are associated with particular mental states (Carlson, Schrater, & He, 2003; Cox & Savoy, 2003; Haxby et al., 2001; Haynes, 2015; Kamitani & Tong, 2005; Kriegeskorte, Goebel, & Bandettini, 2006). This approach targets informational content in patterns of activity across multiple variables, rather than differences in activity in single variables (Kriegeskorte et al., 2006). MVPA is thus a useful tool in the understanding of emotional space - particularly as modern emotion theories suggest emotions to exist in representational space within network-based frameworks, rather than based on isometric, one-to-one relationships between areas of the brain and individual emotions (Hamann, 2012). The Representational Similarity Analysis (RSA) approach (Kriegeskorte & Kievit, 2013; Kriegeskorte, Mur, & Bandettini, 2008) extends the MVPA “brain decoding” approach by modeling the representational content of information in brain activity patterns. RSA allows comparisons of the structure of information in brain activity patterns to theoretical models of cognition (Kriegeskorte, Mur, & Bandettini, 2008).

In the context of defining the representational space of emotion constructs, several recent studies have applied MVPA to examine spatial regions of the brain associated with specific emotions - typically using functional magnetic resonance imaging (fMRI) or positron emission tomography (PET) techniques with high spatial resolution (but usually low temporal resolution). In a review of recent literature, Kragel and LaBar (2016) suggest that MVPA is able to predict neural activity by both emotional dimensions of valence and arousal and discrete emotions. Evidence for this comes from recent decoding studies which suggest that both emotional dimensions and distinguished emotion categories can be observed using MVPA (e.g., Kragel & Labar, 2013; Kragel & LaBar, 2014; Saarimäki et al., 2016).

Far fewer neuroimaging studies have examined temporal signatures of emotional constructs as a way to determine how different emotions are classified. This is despite the useful information that temporal signatures can provide in understanding the categorization of emotions – particularly as a way to possibly separate brain processes involved in representing multiple emotional constructs (see Waugh, Shing, & Avery, 2015). Researchers who have examined temporal signatures suggest that different, discrete emotions elicit unique temporal neural signatures (Costa et al., 2014; Eger, Jedynak, Iwaki, & Skrandies, 2003; Esslen, Pascual-Marqui, Hell, Kochi, & Lehmann, 2004). For example, in an event-related potential (ERP) study (Costa et al., 2014), participants passively viewed a subset of images from the International Affective Picture System (IAPS; Lang, Bradley, & Cuthbert, 2008). Afterward, participants categorized the images into four emotional categories (fear, disgust, happiness, or sadness) and rated the images on the dimensions of valence and arousal on a 9-point scale. Costa and colleagues found that emotion-specific time signatures differentiated between discrete emotions, with the unique time signature of fear occurring fastest, followed by disgust, then happiness, then sadness. Thus, MVPA reveals that temporal dynamics also differentiate between discrete emotion categories (Costa et al., 2014).

Previous studies examining temporal signatures of emotion constructs have mostly used stimuli predefined based on discrete emotional categories (e.g., face stimuli exhibiting particular discrete expressions, Eger et al., 2003; Esslen et al., 2004; images in predetermined categorized and analyzed as separate discrete emotions, Costa et al., 2014). However, as expressed in several emotion theories, it may not always be so easy to isolate a given emotion in any given stimulus. Some disgusting images, for example, can also induce fear. Some pleasant images elicit more happiness than others. And if dimensional properties determine later emotional categorization, the timeline of such dimensional properties should be tracked separately from the discrete emotional properties. Rather than separating images into *a priori*, categories, we used scaled emotional ratings to define all images with emotional categorical weights along all emotional constructs of interest. By doing so, we could better ensure that differences in neural signature were due to differences in emotional ratings per se, rather than reflecting predetermined categories of discrete emotions.

We sought to understand how the brain quickly characterizes emotional constructs in representational space. We applied RSA to high temporal resolution magneto-encephalography (MEG) data to examine time varying neural activity. Compared to EEG, MEG signal is less smeared over sensors and is less distorted by ocular or muscular artefacts (Baillet, 2017), and it therefore requires fewer trials per condition which is perfectly suited for a condition-rich approach such as RSA (Grootswagers, Wardle, & Carlson, 2017; Kriegeskorte, Mur, & Bandettini, 2008). We measured evoked responses with MEG while participants viewed images and engaged in a one-back task. We employed a one-back task in order to encourage participants to attend to the visual properties of the image rather than the emotional construct per se. The images represented scenes of varying emotional content, and were each previously rated on the constructs of valence, arousal, anger, sadness, disgust, fear, and happiness. Using RSA, we examined how reliably image ratings represented neural patterns across emotional constructs, as a way to better determine how the brain represents them.

## 2. Methods

The current study consisted of two parts. In the first part, we obtained behavioral ratings of emotional responses to our stimulus set. The second part involved presenting the images to a new group of participants while their brain responses were recorded using magneto-encephalography (MEG). The MEG recordings were then analyzed using representational similarity analysis framework (RSA; Kriegeskorte, 2011; Kriegeskorte & Kievit, 2013), where we assessed how much information in the MEG recordings of participants’ brain activity could be explained by the behavioral ratings of emotional responses.

### 2.1. Stimuli

Stimuli consisted of 525 visual images. Some images were chosen from the International Affective Picture System (IAPS; Lang, Bradley, & Cuthbert, 2008), some images were chosen from the Nencki Affective Picture System (NAPS; Marchewka, Żurawski, Jednoróg, & Grabowska, 2014), and all other images were chosen from publicly available internet sources. Images were colored, depicted natural scenes, and were chosen to represent different emotional categories: fear (e.g., threatening animals, gunmen, attackers), sadness (e.g., individuals crying, malnourished children, scenes of injustice), erotica, extreme sports, disgust (e.g., medical injuries, rotten food, roadkill), pleasantness (e.g., laughing babies, fuzzy baby animals), and neutral images (e.g., portraits with neutral expressions, individuals playing chess). These categories were only used as a way to ensure a variety of different emotional images – participant ratings were used to determine the emotionality of individual images.

### 2.2. Behavioral ratings

399 participants were recruited on Amazons Mechanical Turk for a ~30-minute task.^1^ Age and sex of participants were not recorded, but Mechanical Turk participants have been found to be representative of the general US population (Berinsky, Huber, & Lenz, 2012). The ratings task was programmed and run on Qualtrics, and participants were randomly assigned to one of eleven picture set conditions. Picture sets contained either 49 or 42 images, and each set contained an equal amount of exemplars (6 or 7) for the categories of fear, sadness, erotica, extreme sports, disgust, pleasantness, and neutral images. An individual picture set always contained the same images, but the images were presented in a different, random order for every participant in that condition. Across all sets, all 525 images were rated. For each image, participants gave their response on a scale of 1-9 for the following seven questions:

1. How HAPPY/UNHAPPY does this picture make you feel (on a scale from / 1-very unhappy, to 9-very happy)? (Valence)
2. How CALM/EXCITED does this picture make you feel (on a scale from / 1-not at all arousing to 9-very arousing)? (Arousal)
3. How much does this image make you feel Anger (on a scale from 1-not at / all to 9-very much)?
4. How much does this image make you feel Sadness (on a scale from 1-not at / all to 9-very much)?
5. How much does this image make you feel Fear (on a scale from 1-not at / all to 9-very much)?
6. How much does this image make you feel Disgust (on a scale from 1-not at / all to 9-very much)?
7. How much does this image make you feel Happiness (on a scale from 1-not at / all to 9- very much)?

At the end of the task, participants were debriefed about the purpose of the ratings task, and were compensated $1.50 through MTurk. Participants gave informed consent through instructions at the beginning of the study, and the experiment was approved by the UNSW Sydney Human Research Ethics Approval Panel. Mean responses for each question of every image were computed over participants, and used as inputs for RSA. These means are available for download at OSF: https://osf.io/5zqfa/.

### 2.3. MEG data acquisition

13 healthy volunteers (9 female, mean age=24.7 years (*SD*=4 years), all right handed, with normal or corrected-to-normal vision) participated in the MEG part of the study. All participants were recruited from the Macquarie University Student Participant pool, and gave written consent prior to the study. Participants were financially compensated for their time. The study was conducted with the approval of the Macquarie University Human Research Ethics Committee.

For each participant, we used a unique combination of 99 stimuli, of which 49 were seen by all participants. The set of stimuli seen by all participants was included to quantify the between subject variance in the responses, by computing the noise ceiling (described below). The sets of unique stimuli were selected to cover stimuli from all stimulus categories (e.g., happy, sad, neutral, etc.) and to have approximately similar behavioral rating distributions. A table with individual participant image set rating means, and a scatterplot of stimuli-wise intensity values of emotional constructs related to every other emotional construct, are available on OSF: https://osf.io/5zqfa/. Before starting the MEG session, to familiarize the participant with the stimuli, they were shown their individual subset of stimuli while rating them on valence, using a 1-9 key response to the question “how happy does this picture make you feel?”. This gave participants an opportunity to look at each image for a lengthy amount of time and identify their emotional content, to allow participants to better identify each image in the scanner and know their emotional content while doing a visually-based task. Next, participants lay supine inside a magnetically shielded room (Fujihara Co. Ltd., Tokyo, Japan) while the MEG signal was sampled at 1000 Hz from 160 axial gradiometers (Model PQ1160R-N2, KIT, Kanazawa, Japan). Recordings were filtered online between 0.03 Hz and 200 Hz. Stimuli were presented in random order for 200 ms each, with an inter-stimulus interval that varied randomly between 850-950ms. Each stimulus was presented 32 times throughout the experiment. Participants were instructed to press a button when a stimulus repeated during the inter-stimulus interval. The repeating stimuli were counterbalanced throughout the experiment. The relatively long temporal distance between images allowed participants to respond to repeated images, and also prevented possible emotion-induced blindness - such that with this delay, emotional images would likely not interfere with the processing of subsequent images (Most, Chun, Widders, & Zald, 2005). Using the Yokogawa MEG Reader Toolbox for MATLAB (Yokogawa Electric Corporation, 2011), we extracted –100 to 600 msec of MEG data relative to the onset of the stimulus in each trial. Next, the data were downsampled to 200Hz, and four trials of each stimulus were averaged to create 8 pseudotrials per stimulus (Grootswagers et al., 2017; Isik, Meyers, Leibo, & Poggio, 2014). Note that no other preprocessing steps were performed on the data (e.g., no channel selection, baseline correction, artefact removal, etc.).

#### 2.4. Representational similarity analysis

The data were analyzed using RSA (Kriegeskorte, 2011; Kriegeskorte, Mur, Ruff, et al., 2008; Kriegeskorte & Kievit, 2013; Kriegeskorte, Mur, & Bandettini, 2008). This approach works by first creating representational dissimilarity matrices (RDMs) for each subject that describe the difference in the neural response between pairs of stimuli from the recorded brain activity (described below). These neural RDMs can then be compared to ‘candidate’ RDMs, such as behavioral rating RDMs (Redcay & Carlson, 2015; Wardle, Kriegeskorte, Grootswagers, Khaligh-Razavi, & Carlson, 2016), or can be compared to other modalities (Cichy, Pantazis, & Oliva, 2014) and other species (Kriegeskorte, Mur, Ruff, et al., 2008).

We constructed time-varying RDMs for each subject. A 25 msec sliding window approach was used where at each time point (Figure 1A), we included data from the four preceding time points (to avoid artificially moving the onset of information in the data). The channel activations at these five time points were concatenated into a feature vector for each trial. A linear discriminant t-value (LD-t) between the feature vectors of the trials belonging to two stimuli was computed and stored in a NxN matrix (Figure 1B), where N is the number of stimuli. The LD-t is a cross-validated measure of dissimilarity, similar to a cross-validated Mahalanobis distance (Nili et al., 2014; Walther et al., 2015). LD-t values for all possible pairwise combinations of subject-specific stimuli were computed, yielding an NxN representational dissimilarity matrix (RDM, Figure 1C). Repeating the process at each time point resulted in a set of time-varying RDMs for each subject (Figure 1D).

**Figure 1.**
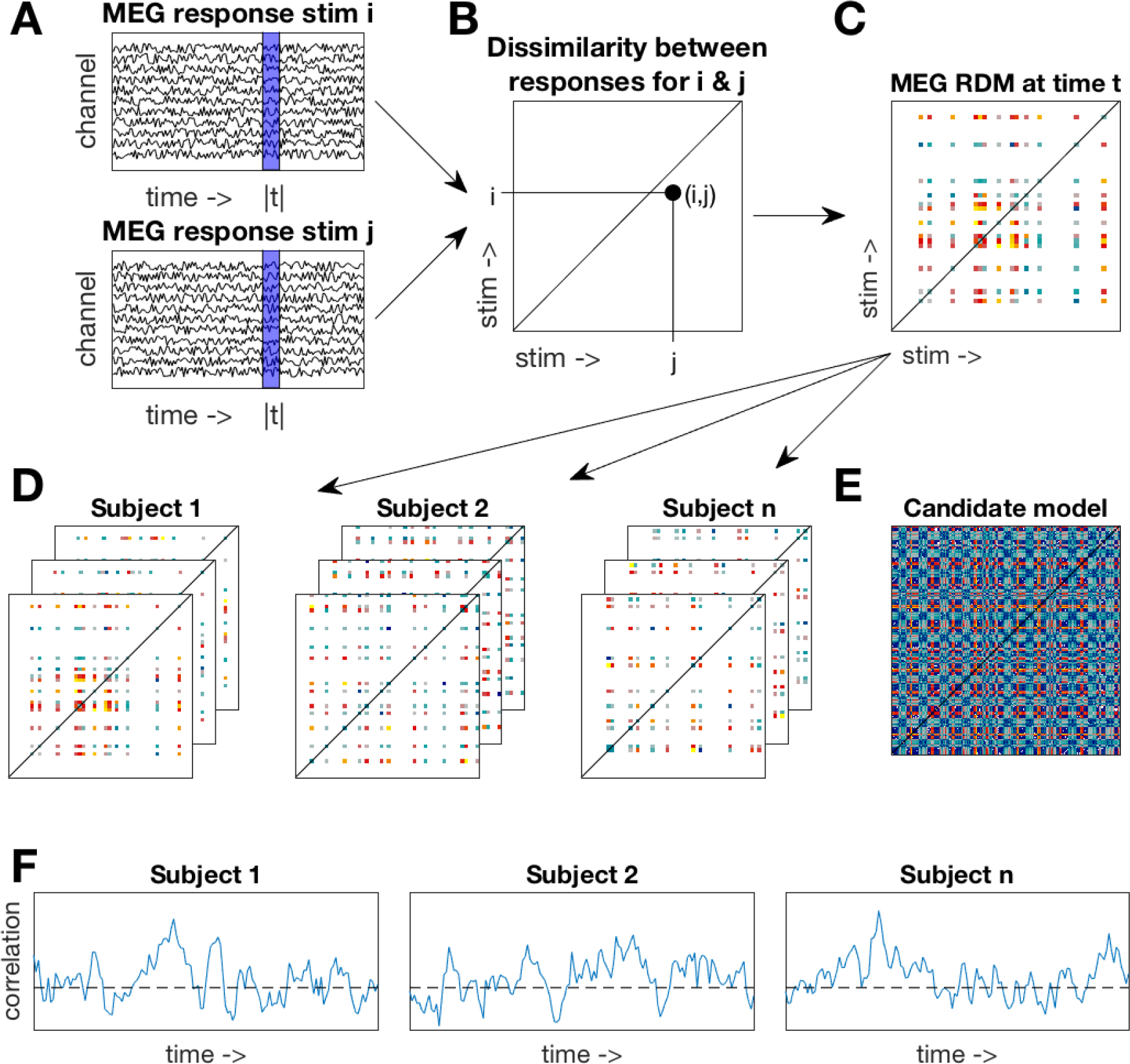
Methods overview. **A.** A 25msec sliding window approach was used. At one time window (blue area), MEG responses to two stimuli were extracted. **B.** A dissimilarity value between MEG responses was computed and stored in an N=N matrix. **C.** Repeating the process over all stimuli yielded a subject-specific representational dissimilarity matrix (RDM). Note that each subject saw a subset of the stimuli, resulting in only a partially filled RDM. **D.** A time-varying RDM was created for each subject by repeating the process over all time points. **E.** One example candidate model RDM. Multiple candidate models were constructed from the behavioral ratings and HMAX model. **F.** For each subject, their time-varying RDM was correlated to the candidate model (restricted to their respective stimulus subset) resulting in a time-varying correlation with the candidate model. The time-varying correlations were assessed for significance at the group level.

Candidate model RDMs were created using the behavioral rating data (e.g., Figure 1E). For each of the rating questions, an RDM was constructed by taking the difference in the mean ratings for all pairs of stimuli. This resulted in seven behavioral rating RDMs. In addition, HMAX (Riesenhuber & Poggio, 2002; Serre, Wolf, & Poggio, 2005) was used as a control model to approximate the visual responses to the images. The first two layers (S1 and C1) of HMAX represent V1 simple and complex cells. The higher layers (S2 and C2) represent complex and invariant features that pool from the lower two layers, in a similar manner to the ventral temporal cortex. The responses to our stimuli of the units in each layer were concatenated into vectors. We then computed pairwise dissimilarities on these vectors to create four RDMs (one per HMAX layer). To assess the overlap of information amongst our candidate RDMs, we computed similarity values (Spearman’s rho) for all pairwise combinations of the RDMs. These were then visualized using multidimensional scaling, which represents the multidimensional RDMs as points in a lower dimensional 2D space, and arranging them so that distance between points approximate the similarity between exemplars in the RDMs.

The 11 candidate model RDMs (4 HMAX RDMs and 7 behavioral RDMs) were compared against the time-varying neural RDMs by computing the correlation (Spearman's rho) between the candidate model RDMs (Figure 1E) and the neural RDMs at each time point (Figure 1D), yielding a time-varying correlation for each candidate RDM for each subject (Figure 1F). For each subject, the candidate models were restricted to the 99 stimuli in that subject’s stimulus set (see Figure 1C,D). The correlations for the behavioral rating models were computed using partial correlations, to control for the correlations with the HMAX C2 RDM (which had the highest correlation with the neural data). In addition, we used partial correlations for assessing to what extent the correlations with the discrete rating models could be explained by the valence and arousal models. Wilcoxon signed rank tests (with subject as random effect) were then used at each time point for statistical inference on the correlations at the group level. False Discovery Rate (FDR) was used to correct for multiple comparisons by determining the FDR-threshold (q = 0.05) and subsequently using this to threshold the p-values.

To estimate the range of correlations to be expected given the between-subject variance, we computed the noise ceilings for our data (as described in Nili et al., 2014). The noise ceiling was computed using only the stimuli that were shared across participants. This means that the true noise ceiling is likely to be lower that our estimate, as including more stimuli (with fewer participants per stimulus) will increase the noise in the data.

### 3. Results

The aim of this study was to study the brain’s time varying representation of different emotional constructs. We used RSA to compare the brain’s time varying representation of the stimuli to candidate models that were created using behavioral ratings on emotional dimensions and intensity of discrete emotions. HMAX was used to create control models of low-level image properties.

To assess the similarities amongst the candidate RDMs, we correlated the pairwise combinations of the RDMs (Figure 2A), and visualized the relations between the RDMs using multidimensional scaling (Figure 2B). These results show that HMAX RDMs cluster together, with the exception of layer C2, which is different from the other layers. The behavioral rating RDMs are different from the HMAX RDMs. Fear and arousal are the most different from the other rating RDMs. We computed the noise ceiling for the MEG RDMs based on the stimuli that were seen by all subjects (Figure 2C). The noise ceiling estimates the range of the maximum possible correlation between the data and any model, given the between-subject variance in the data. For comparison, the correlation between candidate models and the MEG data are displayed with the noise ceiling. The HMAX RDM correlations (blue lines) in the early response reach about 25% of the noise ceiling, and the candidate models (red and yellow lines) in the later response are approximately half as strong as the lower bound on the noise ceiling. Note that the noise ceiling is an estimate of the correlation of the “true model”. The emotional dynamics are likely to be covered using a combination of several candidate models. Therefore, the correlations of the individual models are not expected reach the noise ceiling.

**Figure 2.**
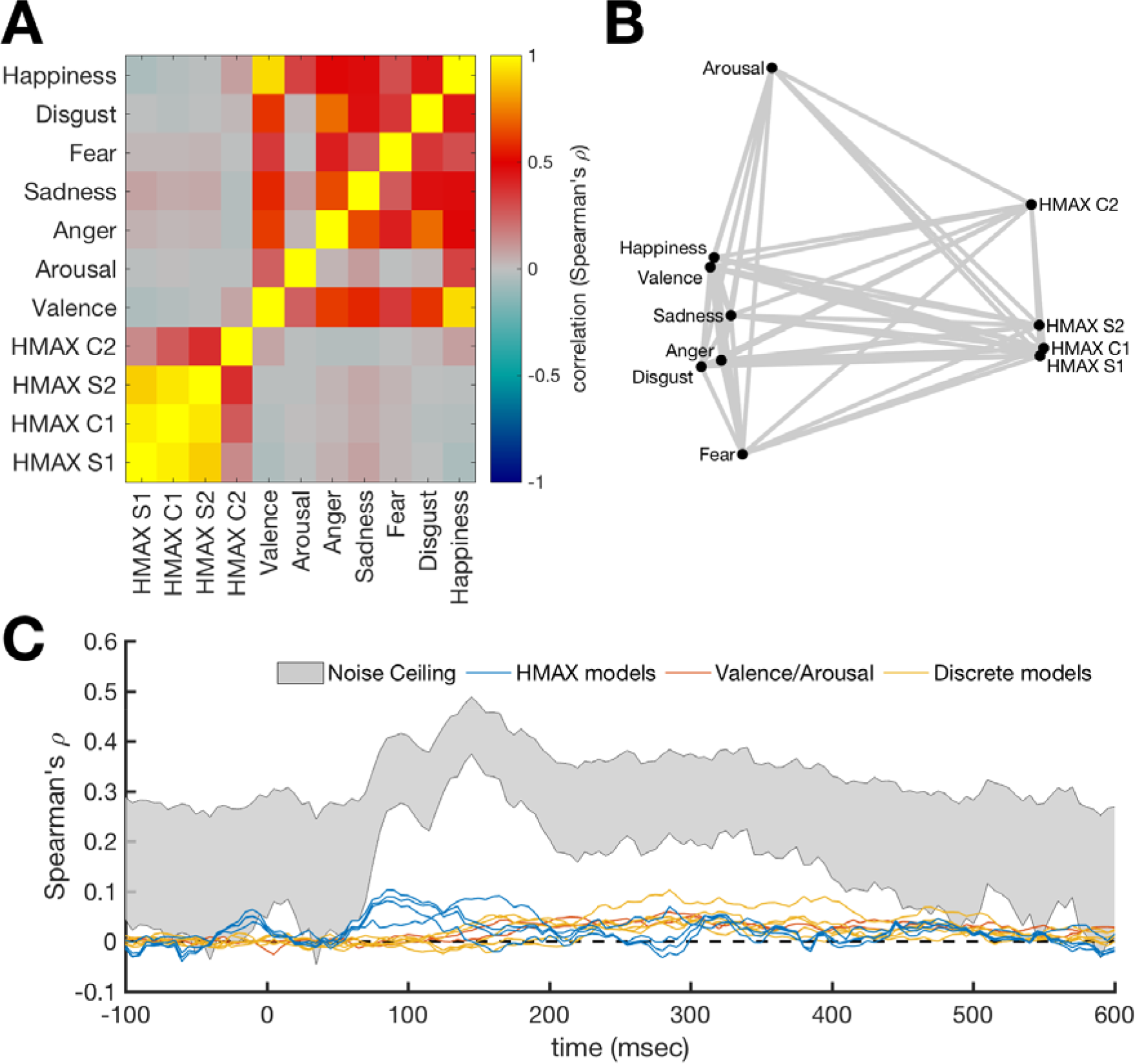
Similarity between candidate RDMs. **A.** Correlation between candidate RDMs. To assess overlap of information between the RDMs, the candidate RDMs were correlated with each other. Higher values mean the RDMs are similar. **B.** Visualization of the similarities between candidate RDMs. Points represent RDMs and distances between these points represents the dissimilarities (1-correlation) between the RDMs. This shows that HMAX RDMs cluster together and are different from the behavioral rating RDMs. **C.** RDM correlations with the MEG time-series RDM compared to noise ceiling estimates. The noise ceiling estimates the range of the maximum correlation that the true model can have with the data, given the between-subject variance. The emotional dynamics are likely to be coveredusing a multifaceted model. Therefore, the correlations of the individual candidate models are not expected reach the noise ceiling.

The time-varying neural RDMs were correlated at each time point with the HMAX RDMs that capture low level visual responses to the images. The correlations for the earliest HMAX layers become significant around 70 msec, which roughly corresponds to visual information entering the striate cortex (Thorpe, Fize, & Marlot, 1996). The correlations with the most complex HMAX layer (C2), which emulates processing areas further along the ventral stream, reach significance later (100 msec), and peaks later (165 msec) compared to the early layers. The C2 layer correlations were significant for the longest time period. The time-varying correlations are shown in Figure 3, where for each HMAX layer, the complete RDM is depicted in the left column. The right column shows their respective time-varying correlations with the neural data. In sum, these results show that the early MEG response to the stimuli is explained well by low-level visual information.

**Figure 3.**
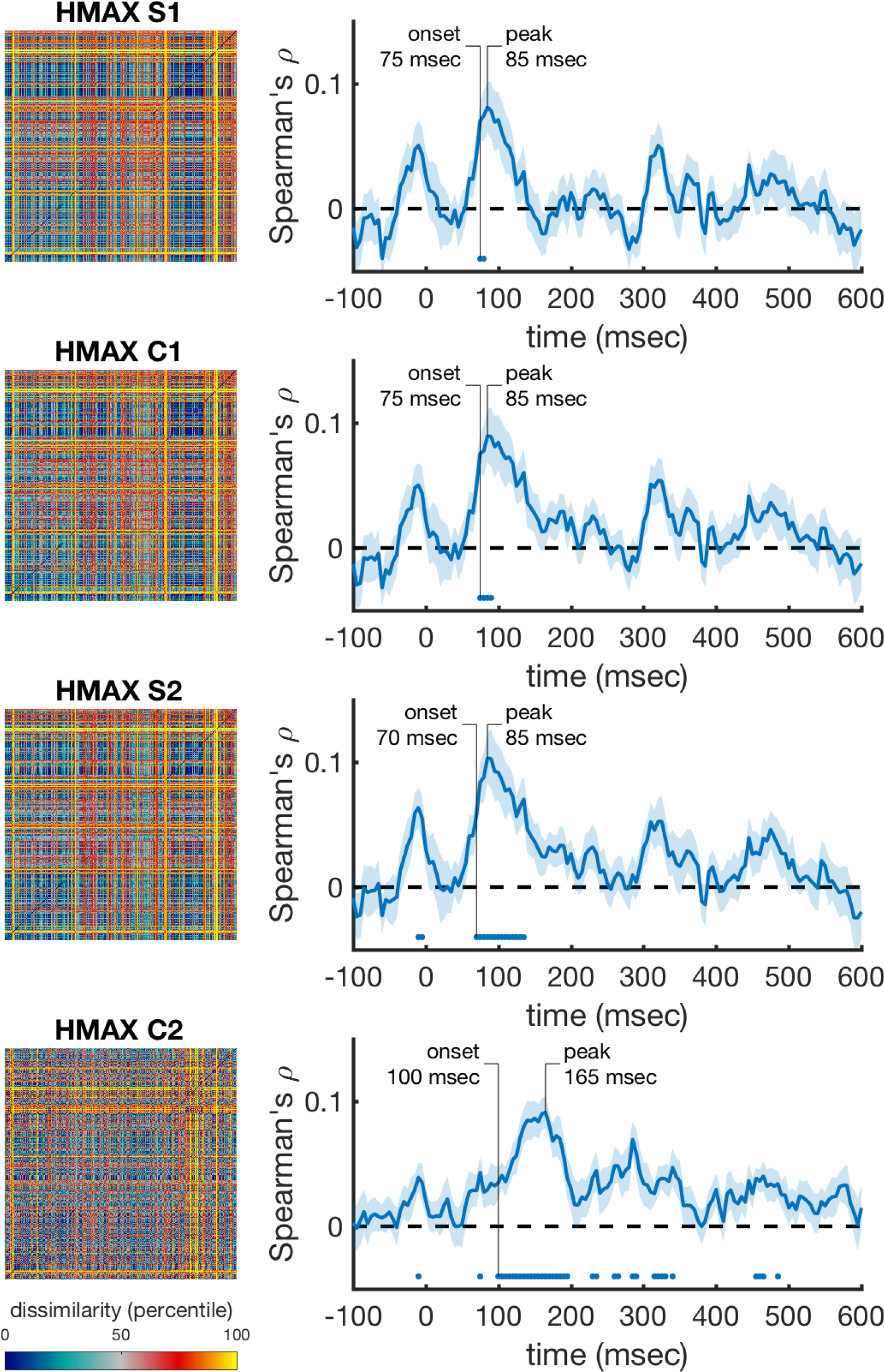
Performance of the low level visual models over time (HMAX layers) The rank-transformed RDMs are shown on the left, and their respective correlations with the time-varying neural RDMs are shown on the right. Marks above the x-axis show significant above-zero correlations (p<0.05 fdr-corrected). Shaded areas represent standard error over participants. Peak correlation and the onset of a sustained significant correlation are annotated in each trace.

Next, we correlated the behavioral rating RDMs with the time-varying neural RDMs, while controlling for the information that is captured by the best performing visual control model, the HMAX C2 RDM (using a partial correlation). We found that valence and arousal are significantly represented in the neural data starting at 145 msec and 175 msec and peaking at 285 msec and 300 msec. Figure 4 shows these correlations, with the rating RDMs in the left column, and their respective correlation with the neural data in the right column. Discrete emotion categories were represented early as well (Figure 5). Correlations with anger ratings started relatively late (300 msec) in the time series (Figure 5A). The correlations for sadness did not reach significance, but were consistently above zero from after around 240 msec onward (Figure 5B). The onset for fear was the earliest of all, starting at 140 msec, and had an early (175 msec) peak (Figure 5C). Disgust showed the largest and most sustained significant correlation over time, starting at 170 msec and peaking at 285 msec (Figure 5D). Happiness showed a lower, but fast (145 msec) and sustained significant correlation over time (Figure 5E).

**Figure 4.**
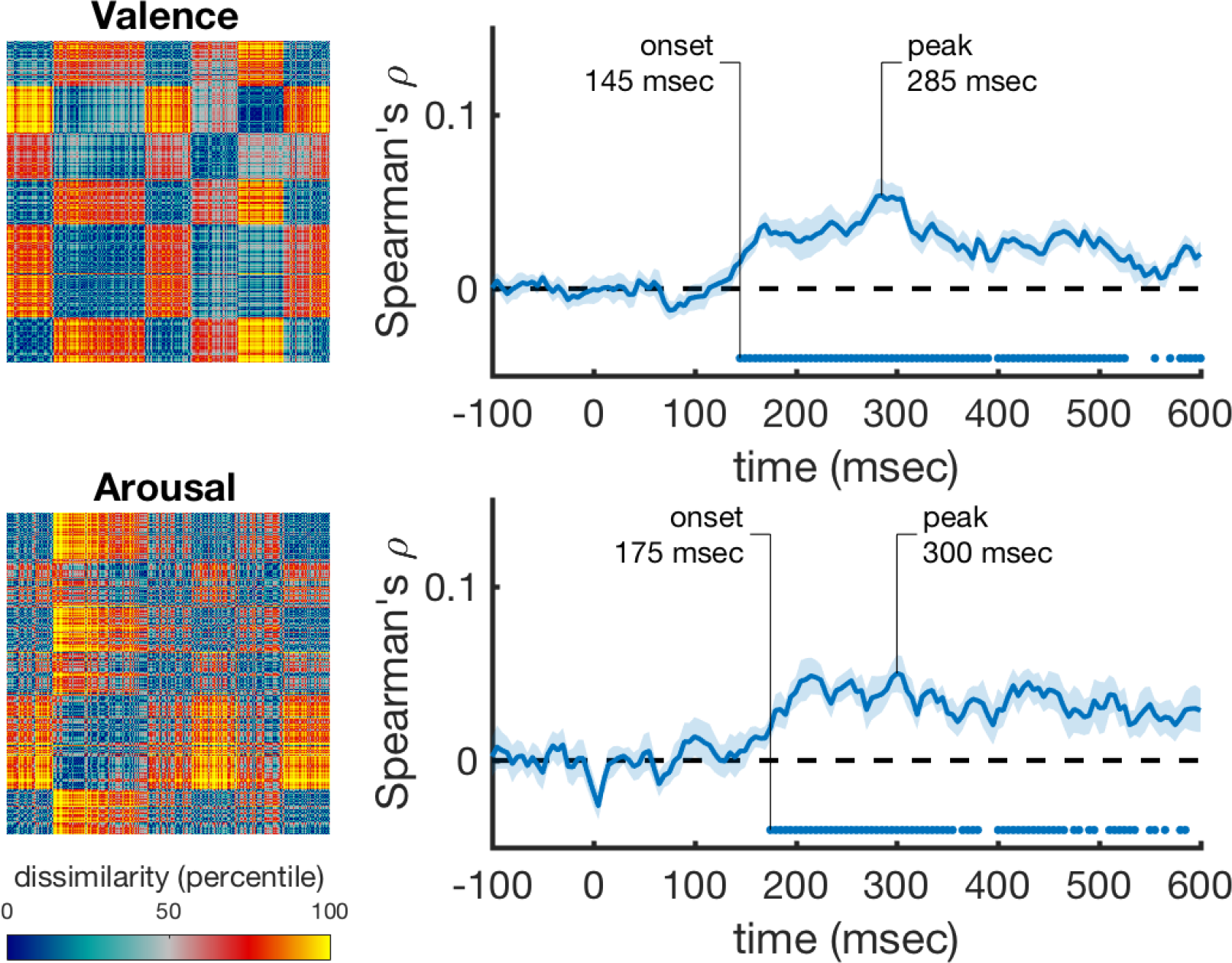
Performance of the valence and arousal rating scale models over time after correcting for HMAX correlations. The rank-transformed RDMs are shown on the left, and their respective correlations with the time-varying neural RDMs are shown on the right. Correlations were computed using partial correlations to control for the correlations with the HMAX C2 model. Marks above the x-axis show significant above-zero correlations (p<0.05 fdr-corrected). Shaded areas represent standard error over participants. Peak correlation and the onset of a sustained significant correlation are annotated in each trace.

**Figure 5.**
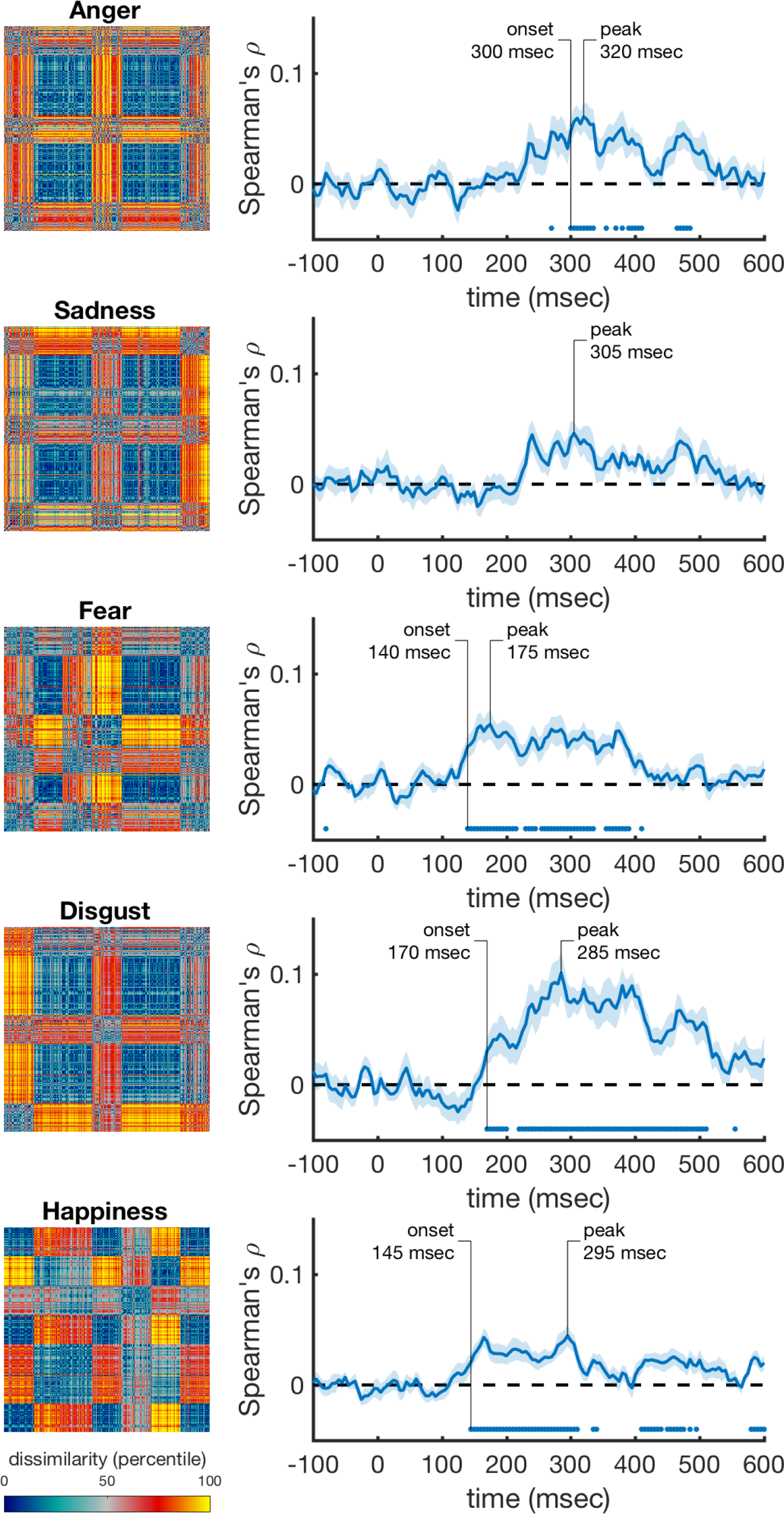
Performance of the discrete emotion rating scale models over time after correcting for HMAX correlations. The rank-transformed RDMs are shown on the left, and their respective correlations with the time-varying neural RDMs are shown on the right. Correlations were computed using partial correlations to control for the correlations with the HMAX C2 model. Marks above the x-axis show significant above-zero correlations (p<0.05 fdr-corrected). Shaded areas represent standard error over participants. Peak correlation and the onset of a sustained significant correlation are annotated in each trace.

We then asked whether the correlations of the discrete emotion categories explain different aspects of the representation than the correlations for valence and arousal. We correlated the discrete emotion rating models while controlling for the effects of HMAX, valence, and arousal. The fear and disgust rating models still correlated significantly with the data (Figure 6), and thus explain variance that is not captured by the valence and arousal models. However, correlations with all other rating models were not significant after correcting for valence and arousal.

**Figure 6.**
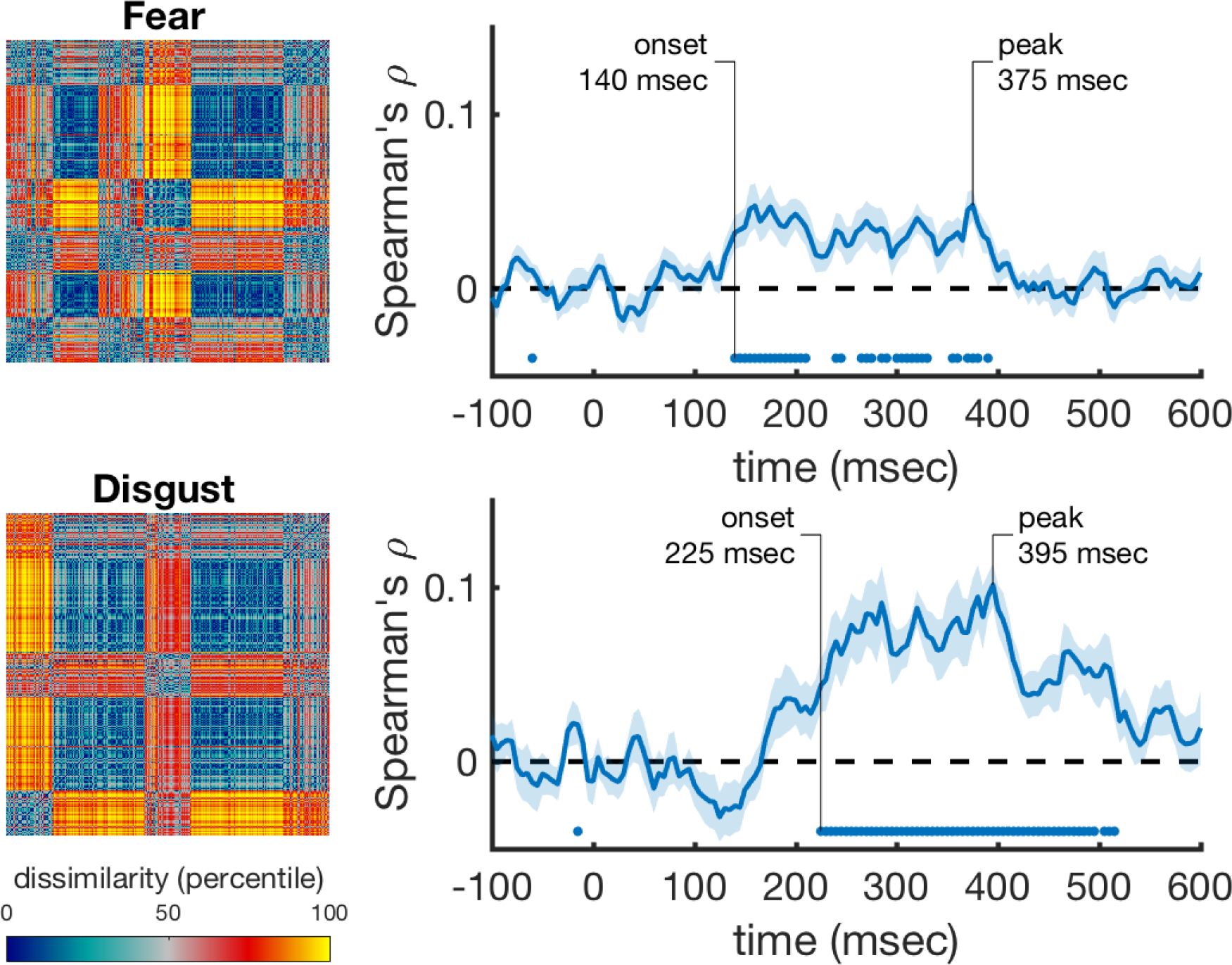
Performance of the fear and disgust rating scale models over time after correcting for HMAX and valence/arousal correlations. The rank-transformed RDMs are shown on the left, and their respective correlations with the time-varying neural RDMs are shown on the right. Correlations were computed using partial correlations to control for the correlations with the HMAX C2 model, and with the valence and arousal models. Marks above the x-axis show significant above-zero correlations (p<0.05 fdr-corrected). Shaded areas represent standard error over participants. Peak correlation and the onset of a sustained significant correlation are annotated in each trace.

## 4. Discussion

We examined early neural signatures associated with different emotional constructs, as a way to explore how the brain characterizes the representational space of emotion. To model the early neural correlates of emotional constructs, participants viewed behaviorally rated images inside a MEG scanner, and we applied RSA to examine the time course of representation along emotional dimensions of valence and arousal, and along discrete emotional categories of sadness, fear, disgust, sadness, and happiness. Our data suggest that both arousal and valence were represented relatively early in the brain. This was despite the range of image types that we had in our study of varying arousal and valence levels. The discrete emotions of fear, disgust, and happiness also had reliable, early signatures, anger was reliably represented at a later processing stage, and sadness was not reliably represented during our measurement window. Moreover, despite early representations in both disgust and fear (both classified as negatively valenced and highly arousing emotions), the neural signature of disgust lasted for a longer amount of time than that to fear. Disgust also had a distinct signature, such that its temporal signature showed a higher correlation than most other emotion properties at its maximum peak. While speculative, this result may suggest that disgust in particular elicits a strong response, and that there may be more similarity in people’s disgust response than their response to other emotional categories.

In the context of traditional emotion classification theories – which suggest emotional space exists either along dimensions (e.g., Russell, 1980) or discrete emotional constructs with distinct qualities (e.g., Ekman, 1992) – these data suggest differences in neural temporal representational signature across at least some discrete emotion constructs in addition to the basic dimensional qualities of valence and arousal. Thus, in early neural processing, emotions are differentiable by RSA. In particular, distinct fear and disgust representations were observed when correcting for valence and arousal ratings, suggesting that these discrete emotions can be observed above and beyond valence and arousal.

In the framework of constructivist theories of emotion (Barrett, 2017; Cunningham et al., 2014; Lindquist et al., 2013), our finding that valence and arousal dimensions predict early temporal patterns of activation supports the notion that dimensional affect plays a role in how we assign emotional meaning. Additionally, the different time courses linked with discrete emotion categories may be suggestive of the relative rapidity with which we may categorize and construct our experience of different discrete emotions. Constructivist theories suggest that “core affect” (based on dimensional constructs like valence and arousal) is the basis with which we extract the experience of discrete emotions (Barrett, 2017). Mostly consistent with this account, the predictiveness of valence and arousal had rapid onset, with ratings of discrete emotions occurring later - with one exception. Ratings of fear had predictive value as early than ratings of valence and arousal. It may be that fear (due to its biological importance) is fast-tracked in processing unlike other dimensional emotions (e.g., LeDoux, 2000; Morris et al., 1999; Tamietto & de Gelder, 2010). Moreover, representations of fear and disgust were observed when we corrected for valence and arousal correlations, to suggest that these emotions in particular may recruit from different processes. Altogether, these results suggest distinct neural temporal signatures of different emotion categories, and while speculative, may represent the constructive nature of emotional experience. Future work can examine the different processes for fear and disgust more directly, by using image sets with varying levels of fear and disgust, while matching their distributions of valence and arousal.

To our knowledge, only one other study used MVPA on temporal neural signatures in response to natural scene images of different emotional constructs (Costa et al., 2014). Costa et al. (2014) reported different time signatures for fear and disgust, but found that the onsets of neural responses distinguished them, whereas we found similar onsets but prolonged representation for disgust. Additionally, while Costa et al. reported a slow onset time for happiness and sadness, we found a relatively fast response to happiness, and no significant unique response to sadness at all. The differences in results could be due to the nature of task-demands – participants in our study were actively engaged in a one-back task, whereas participants in Costa et al.’s task were passively viewing the images. Differences could also stem from the way in which emotional stimuli were categorized – Costa et al. defined images into discrete categories, whereas we used scaled ratings to define their categorical weights. Altogether, we similarly conclude that at least some discrete emotions demonstrate unique neural signatures, and that valence and arousal cannot fully explain differences between emotional responses (particularly in fear and disgust).The use of emotional images is common practice in emotion research. Large datasets with hundreds of previously rated images (e.g., IAPS, Lang, Bradley, & Cuthbert, 2008; NAPS, Marchewka, Żurawski, Jednoróg, & Grabowska, 2014) are commonly used to elicit emotional responses in many varied and diverse emotional paradigms. The time signatures of different emotional constructs should be considered in these tasks and the interpretation of results. For example, images depicting anger may take a greater amount of time to be represented compared to fear images. The intensity of emotions to those images may therefore vary depending on when the response is measured.

The type of emotional processing elicited by emotional images should also be considered. In this study, participants viewed images that were themselves emotional, rather than participants inducing emotions organically through imagery, narratives, etc. It is therefore worth noting that emotional constructs in our study were based on the categorical ratings of emotional stimuli rather than the subjective experience of emotion per se. We also only used the dimensions of valence and arousal to represent the traditional dimensional space of emotions, whereas future work may want to incorporate additional dimensions to represent the space with higher dimensionality (see Fontaine et al., 2007). Moreover, previous research indicates specific, modality-dependent activation to represent valence across the modalities of vision and taste (in ventral temporal cortex and anterior insular cortex), as well as modality-independent activation to represent valence (in the lateral orbitofrontal cortices; OFC) (Chikazoe, Lee, Kriegeskorte, & Anderson, 2014). These results indicate there may be modality-specific lower-level representations of valence, but higher-level representations of valence that extend across modalities. Future research should continue to examine how the representation of different types of subjective emotional experience and emotion from different modalities compare with those when viewing intrinsically emotionally powerful stimuli. Another consideration is that we used a one-back task, which involves working memory processes. While the working memory demands were the same across all emotional constructs, it is unclear if the results may differ if images are presented under passive viewing rather than in a one-back task.

Modern imaging and statistical techniques allow us to explore long-debated questions with a new lens. MVP A has particular strengths in the debate of emotional representation, since evidence seems to suggest network organization of emotions, rather than simple isomorphic activity in unique brain structures. Our data suggest that many emotions are distinguishable early in the evoked response, and that at least some emotional constructs carry unique signatures– perhaps reflecting the timeline of the constructive nature of discrete emotions. While these data utilize and represent information that we are able to extract and measure, it still remains unknown what information the brain uses to interpret emotions (cf. De-Wit, Alexander, Ekroll, & Wagemans, 2016). Another open question is which brain areas are involved in the temporal representation of emotion categories. Here, whole-brain MEG was used in the analysis, which does not allow differentiating between the contributions of different brain areas. Future studies could investigate the spatio-temporal dynamics of emotion categories by using our approach on reconstructed activity from different brain areas (using e.g., beamformer techniques (Van Veen, Drongelen, Yuchtman, & Suzuki, 1997)). Nevertheless, since we are able to differentiate signal from both valence and arousal ratings and several distinct emotional categories, in a timeline that might represent the constructive nature of discrete emotion experience, these findings not only inform how the brain may represent emotional constructs, but also shed light on the way that we should discuss, explore, theorize, and ultimately define them.

## Acknowledgements

This work was supported by Australian Research Council (ARC) Future Fellowship [FT120100707] to SBM, and ARC Future Fellowship [FT120100816] and ARC Discovery project [DP160101300] to TAC. We also wish to thank the ARC Centre of Excellence in Cognition and its Disorders (CCD) for their support on this project. The authors declare no conflicts of interest.

## Footnotes

1 Most but not all participants completed the entire ratings task (302 out of 399 participants completed the whole task). To compute average emotion scores for each image, we included ratings regardless of the number of trials participants completed.

